# Discovery of ARQ-501 as a potent PafA inhibitor that elevates KatG levels to potentiate isoniazid in mycobacteria

**DOI:** 10.64898/2026.06.03.729796

**Authors:** Cong Li, Bingzhou Huang, Hang Xiong, Qiaohua Yan, Yan Liu, Cuiting Fang, Youfu Luo, Peng Xu, Tao Luo, Qingxiang Sun

**Affiliations:** Cancer Center and State Key Laboratory of Biotherapy, West China Hospital, Sichuan University, Chengdu 610041, China; Department of Pulmonary and Critical Care Medicine, Sichuan Provincial People’s Hospital, University of Electronic Science and Technology of China, Chengdu 610032, China; Department of Pathogen Biology, West China School of Basic Medical Sciences & Forensic Medicine, Sichuan University, Chengdu 610041, China; State Key Laboratory of Respiratory Disease, Guangzhou Institutes of Biomedicine and Health, Chinese Academy of Sciences, Guangzhou 510530, China; National Clinical Research Center for Infectious Diseases, Shenzhen Clinical Research Center for Tuberculosis, Shenzhen Third People’s Hospital, Shenzhen 518112, China

**Author notes:** Corresponding authors: Peng Xu, Tao Luo, Qingxiang Sun. These authors contributed equally to this work (Cong Li, Bingzhou Huang, Hang Xiong, Qiaohua Yan).

**Keywords:** *Mycobacterium tuberculosis*, PafA inhibitor, pupylation, KatG, isoniazid, drug resistance

## Abstract

The prokaryotic ubiquitin-like protein (Pup) conjugation system (PPS), which is essential for *Mycobacterium tuberculosis* (Mtb) virulence but absent in humans, presents an attractive drug target. Here, we report the discovery of ARQ-501, a quinone-based, covalent, substrate-competitive inhibitor of the Pup ligase PafA. ARQ-501 exhibited potent anti-mycobacterial activity against Mtb under host-mimicking stress conditions and within macrophages. We further identified the catalase-peroxidase KatG, essential for activation of the frontline prodrug isoniazid (INH), as a pupylation substrate. ARQ-501 inhibits KatG pupylation, causing its accumulation and creating a selective synergy with INH. This ‘quantity over quality’ mechanism successfully rescued INH activation by the clinically prevalent KatG S315T mutant in enzymatic assays and enhanced INH efficacy against clinical S315T isolates to variable degrees. This work identifies a novel class of PafA inhibitors and a previously unrecognized role of pupylation in regulating KatG, offering a potential therapeutic avenue to combat drug-resistant tuberculosis.

## Introduction

Tuberculosis (TB) remains the world’s leading infectious cause of death, with drug resistance posing a critical threat to global health.^1^ Isoniazid (INH) is a cornerstone of first-line TB therapy, but resistance to this prodrug is increasingly common. KatG is the enzyme that activates the prodrug INH, and mutations in KatG are the primary cause of INH resistance in *Mycobacterium tuberculosis* (Mtb).^2, 3^ For example, the S315T mutation reduces catalytic activity by approximately 90% and is found in 50–90% of INH-resistant clinical isolates.^4, 5^ This underscores the urgent need for alternative therapeutic strategies.^6^

A key component of Mtb’s virulence is its unique ubiquitin-like pupylation system,^7, 8^ where the ligase PafA mediates protein degradation by conjugating a Pup polypeptide to target proteins.^9^ PafA functions by simultaneously binding ATP, its protein substrate, and Pup, leveraging ATP hydrolysis to catalyze Pup conjugation.^10, 11^ PafA is absent in humans, making it an attractive therapeutic target.^12^ Current PafA inhibitors include AEBSF,^13^ bithionol,^14^ ST1926,^14^ and Pi-2-26.^15^ Among them, AEBSF, ST1926, and Pi-2-26 bind to the Pup binding pocket, whereas bithionol targets the ATP binding pocket. Like Pup-proteasome system (PPS) deletion, Pi-2-26 inhibits Mtb survival under nitric oxide stress.^16, 17^ However, the inhibition potency of these inhibitors is limited and there is a need to develop more potent PafA inhibitors.

In this study, we optimized a high-throughput screening assay to discover two more potent PafA inhibitors, ARQ-501 (hereafter referred to as ARQ) and SF1670 (SF). ARQ and SF are reported anti-cancer agents that target NQO1 and PTEN, respectively.^18, 19^ These compounds covalently bind to a site that impairs the conjugation of different PafA substrates to varying degrees. ARQ, but not SF, inhibits pupylation in mycobacteria, leading to potent *Mtb* killing under acidic reactive nitrogen species conditions and within macrophages. Furthermore, ARQ blocked PafA-mediated pupylation of KatG, enhancing INH activation and thereby synergizing with this first-line anti-TB drug.

## Results

### Identification of ARQ-501 as a potent, covalent, and substrate-competitive PafA inhibitor

From a screen of ∼1,400 compounds at 20 μM, we previously identified ST1926 and bithionol, which inhibited PafA with IC_50_ values of approximately 15 μM. To obtain more potent inhibitors, we screened a larger compound library at a lower compound concentration. The assay temperature was adjusted to 37 °*C* to enhance the signal-to-noise ratio (Fig. S1). After screening ∼5400 compounds from the expanded library, we identified 18 compounds that inhibited PafA activity in plate-based assays by more than 60% at 5 μM concentration (Fig. 1A). The reproducibility of this assay was demonstrated by a second screen with three biological repeats (Fig. 1B). The 11 hits that exhibited more than 70% inhibition of PafA in the second screen were validated in in-solution pupylation assays. At a concentration of 2 μM, several compounds notably inhibited PafA, with compound 17 (ARQ-501) and compound 18 (SF1670) standing out as the strongest inhibitors (Fig. S2). At a concentration of 1 μM whereby the previously identified inhibitors bithionol and ST1926 were ineffective, these two compounds still exhibited strong inhibition of PafA activity (Fig. 1C). Notably, ARQ-501 (ARQ) and SF1670 (SF) both have quinone moieties in their chemical structures (Fig. 1D). The enzymatic IC_50_ values for ARQ and SF were determined to be 0.42 μM and 0.44 μM, respectively (Fig. 1E and S3A).

**Fig. 1.**
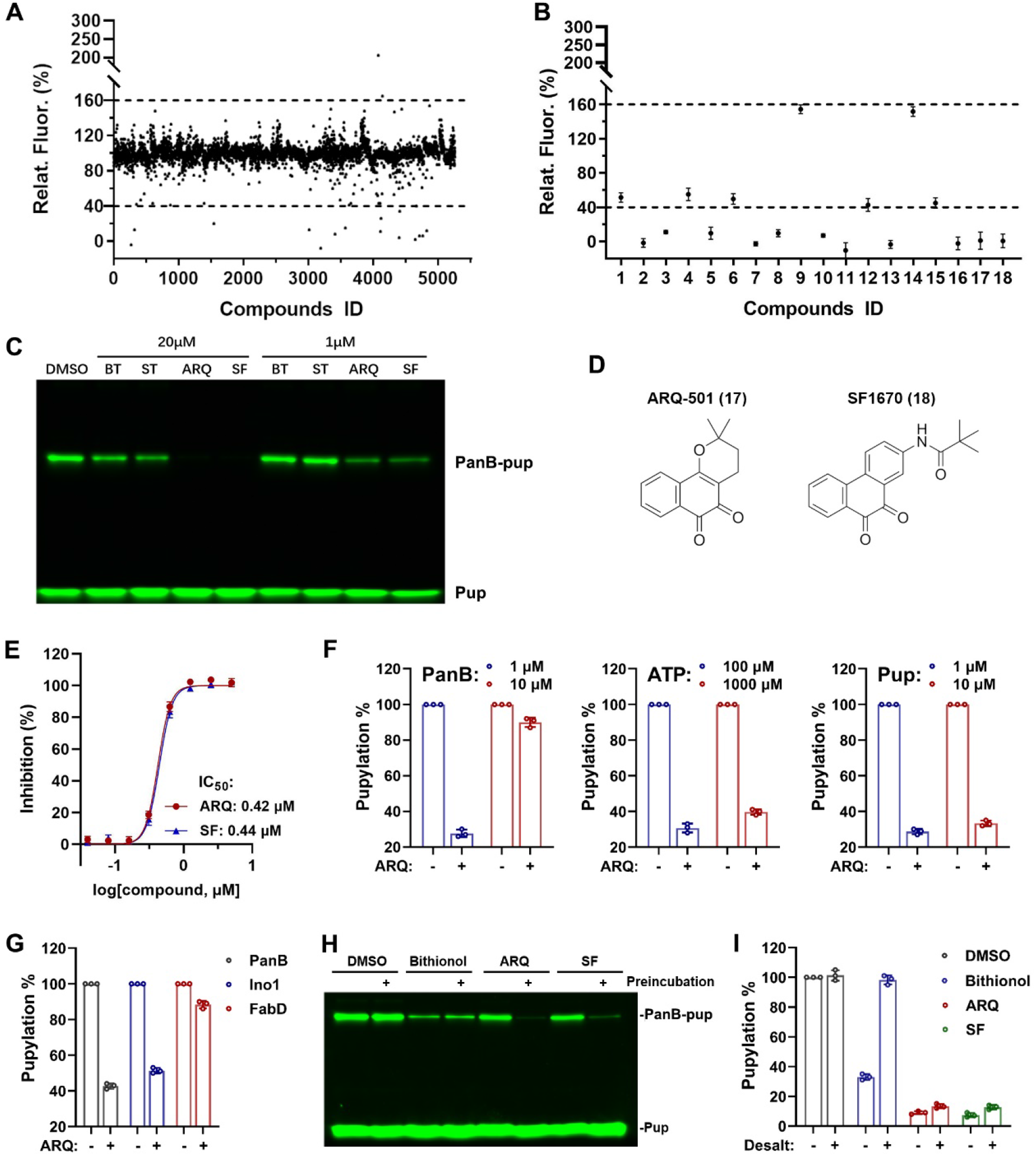
Identification of ARQ-501 as a potent, covalent, and substrate-competitive PafA inhibitor. **(A)** A scatter plot showing the DMSO-normalized relative fluorescence (%) of various compounds during the initial screening. 100% corresponds to DMSO control (no inhibitor). Horizontal dashed lines mark thresholds for significant inhibition or stimulation. **(B)** The second screening on the 18 hits from the first screen with three biological repeats. **(C)** Fluorescent SDS-PAGE image comparing the activity of different compounds at 20 μM and 1 μM concentrations. BT, bithionol; ST, ST1926. **(D)** Chemical structures of compound 17 (ARQ-501) and compound 18 (SF1670). **(E)** Concentration-dependent pupylation inhibition with four-parameter non-linear regression fitting of IC_50_ for ARQ-501 (ARQ) and SF1670 (SF). **(F)** Pupylation changes with varying concentrations of PanB, ATP, and Pup in the presence or absence of ARQ (1 μM). **(G)** Pupylation inhibition of different substrates (PanB, Ino1, and FabD) by ARQ (1 μM). **(H)** PanB pupylation in the presence of bithionol (20 μM), ARQ (1 μM), or SF (1 μM), with or without preincubating PafA (0.2 μM) with inhibitors (25 °C for 2 h). **(I)** Pupylation of PanB using PafA preincubated with bithionol (20 μM), ARQ (1 μM), or SF (1 μM), with or without desalting to remove unbound inhibitors. Data show mean ± standard deviations (SD) from three biological replicates.

To explore the inhibition mechanism, we systematically varied the concentrations of substrates PanB, ATP, and Pup individually to identify the key factor influencing inhibitor activity. The results were unambiguous: increasing the PanB concentration largely abrogated the activity of ARQ and SF, while elevating the levels of ATP and Pup had minimal impact (Fig. 1F, S3B, and S4). This suggests that these inhibitors compete with PanB for the substrate-binding site of PafA. Furthermore, we observed that the same concentration of ARQ inhibited different substrates with varying potencies, reducing PanB pupylation by 55% but FabD pupylation by only 12% (Fig. 1G and S3C). This differential efficacy implies that ARQ likely binds at the periphery of the core substrate-binding pocket, a site that is not equally accessed by all substrates.

We next investigated whether ARQ and SF are covalent inhibitors. The results revealed that their inhibition against PafA was markedly enhanced when preincubated with the enzyme (Fig. 1H). This is in contrast to the noncovalent inhibitor bithionol, which showed comparable activity with or without preincubation (Fig. 1H). As an alternative approach, we incubated PafA with different inhibitors, removed the free inhibitors using a desalting column, and tested the enzyme activity of the treated protein. While the noncovalent reversible inhibitor bithionol lost inhibitory activity upon desalting, ARQ and SF remained active post-treatment (Fig. 1I and S3D), suggesting their covalent engagement with PafA. These results collectively demonstrate that ARQ and SF form covalent bonds with residues at the edge of PafA’s core substrate-binding site.

### ARQ but not SF inhibits pupylation in *Mycobacterium smegmatis*

To assess whether ARQ and SF inhibit pupylation in mycobacteria, we transformed *Mycobacterium smegmatis* (Msm), as a safe surrogate of Mtb, with a plasmid encoding HA-Pup and quantified pupylation levels in the presence of different inhibitors using an anti-HA antibody. At 10 μM, ARQ strongly inhibited pupylation (73.8%), whereas BT produced only a modest inhibitory effect (22.1%) (Fig. 2A,B). In contrast to ARQ, SF was inactive. Although the underlying mechanism remains unclear, this lack of activity may be due to poor cell-wall penetration, metabolic instability, or off-target binding. Further analysis revealed that ARQ inhibited pupylation in Msm in a concentration-dependent manner, with an IC_50_ value of 4.2 μM (Fig. 2C and 2D). Collectively, these results demonstrate that ARQ, but not SF, inhibits pupylation in Msm.

**Fig. 2.**
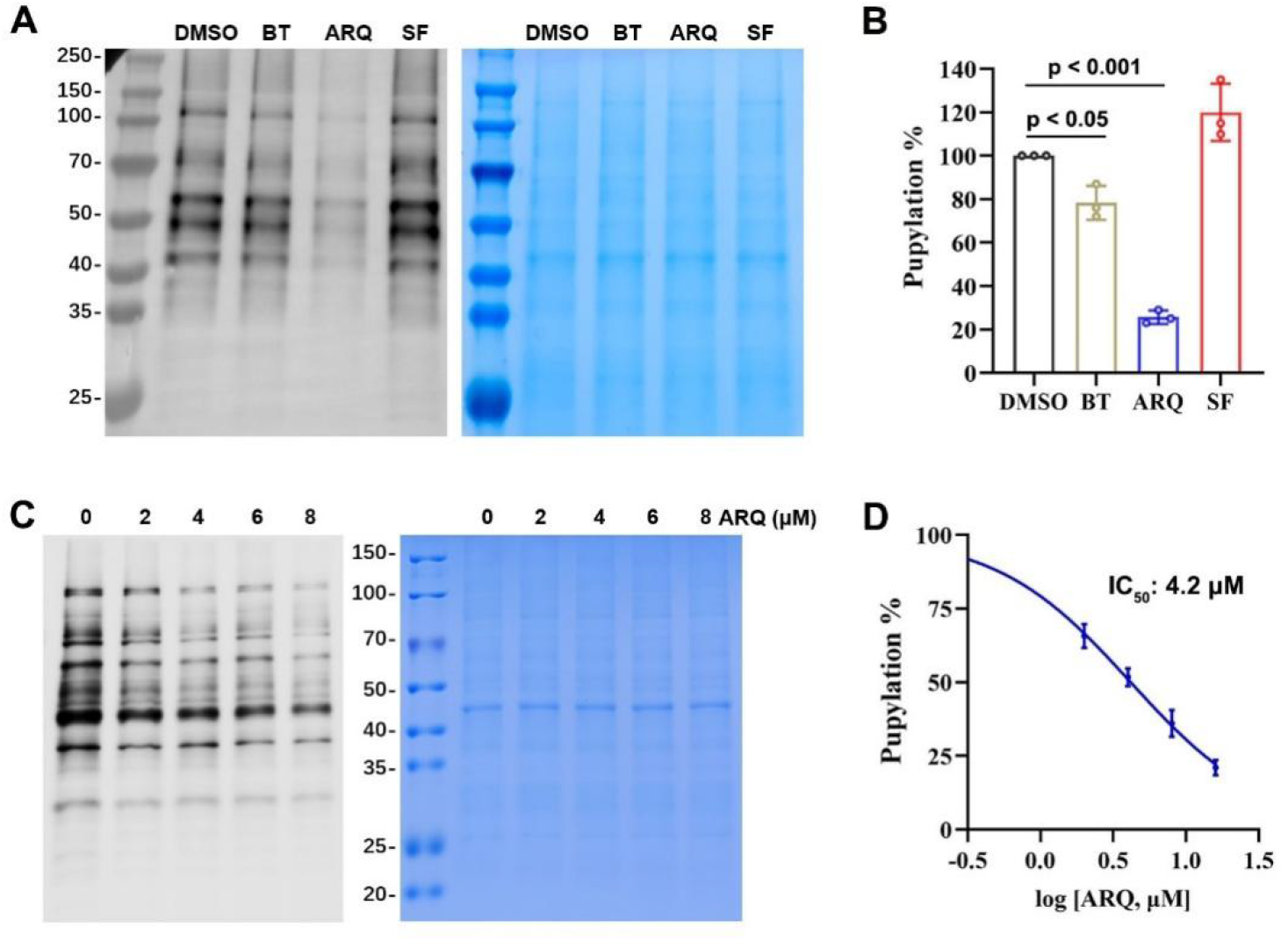
ARQ but not SF inhibits pupylation in Msm. **(A)** Pupylation inhibition in Msm by DMSO (control), BT, ARQ, and SF at a 10 μM concentration. Msm was cultured to OD 0.6, induced with 0.6% acetamide and added with different compounds for three days. The left panel shows the transfected HA-Pup detected by anti-HA antibody, and the right panel is the Coomassie-Blue stained gel indicating equal loading. (**B)** Quantification of pupylation levels in panel A, normalized by the respective Coomassie-Blue intensities. One-way ANOVA was used for statistical comparisons. (**C)** Concentration-dependent inhibition of pupylation by ARQ in Msm. (**D)** Quantification and 4-parameter non-linear IC_50_ fitting of the pupylation levels in panel C. Data show mean ± SD from three biological replicates.

### ARQ inhibits Mtb survival under host-like stress and in macrophages

PafA has been shown to confer mycobacterial resistance to the acidic and reactive nitrogen species (RNS) conditions encountered within macrophages.^17^ Here, we evaluated the impact of ARQ, BT, and SF on Msm growth under normal and acidic RNS stress (3 mM NaNO_2_, pH 5.5). Under normal conditions, none of these compounds showed inhibitory activity on Msm growth at a 10 μM concentration (Fig. 3A). In contrast, in the presence of acidic RNS, ARQ largely abolished Msm growth, whereas BT exhibited moderate inhibition (< 40%) and SF was ineffective (Fig. 3A), aligning with their respective abilities to inhibit pupylation. Dose-dependent growth curves showed that acidic RNS conditions decreased the MIC_90_ of ARQ against Msm by ∼ 80-fold (Fig. 3B), supporting the critical role of pupylation in mycobacterial survival under macrophage-mimicking conditions.

**Fig. 3.**
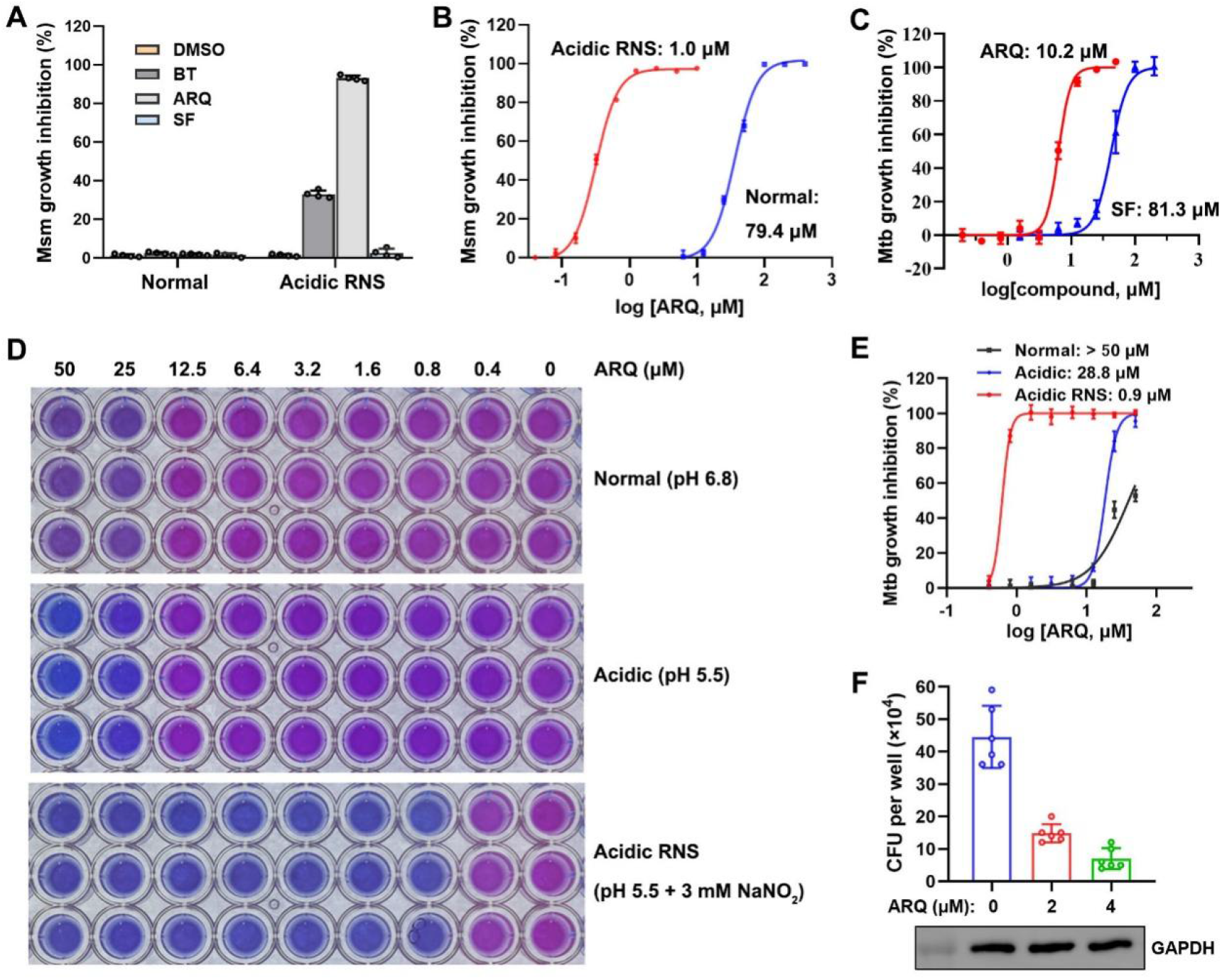
ARQ inhibits Mtb survival in vitro and in macrophages. **(A)** Growth inhibition of Msm by BT (10 μM), ARQ (10 μM), or SF (10 μM) under normal or acidic RNS (pH 5.5 + 3 mM NaNO_2_) conditions. **(B)** Dose-response curve of ARQ on Msm growth under normal or acidic RNS conditions, with MIC_90_ values indicated. **(C)** Dose-response curves of ARQ and SF on Mtb growth inhibition, with MIC_90_ values indicated. **(D)** Dose-response curves of ARQ against Mtb under normal vs. acidic RNS conditions. **(E)** Dose-response curve of ARQ under acidic RNS conditions, with MIC_90_ values indicated. The higher MIC_90_ value for normal condition compared with panel C is due to the higher inoculum (OD_600_ = 0.002 in panel C vs. OD_600_ = 0.2 in this panel). **(F)** Mtb colony forming units (CFU) per well treated with ARQ (0, 2, 4 μM) and the corresponding GAPDH blots confirming the absence of macrophage cytotoxicity. Data are mean ± SD from at least triplicate samples.

Building on these findings in Msm, we assessed ARQ’s efficacy against Mtb H37Rv. Under standard conditions, ARQ displayed an MIC_90_ of 10.2 μM, approximately 8-fold more potent than of SF (Fig. 3C). In the checkerboard assays, acidic RNS stress enhanced ARQ’s potency against Mtb by over 50-fold (Fig. 3D and 3E), mirroring its behavior in Msm. In contrast, acidic RNS only barely enhanced the efficacy of the RNA synthesis inhibitor rifampicin (Fig. S5, from 0.17 μM to 0.14 μM), indicating that the stress-dependent potentiation is specific to ARQ rather than a general effect on antibacterial agents.

We further investigated ARQ’s efficacy in reducing Mtb burden within macrophages. Notably, ARQ reduced intracellular Mtb burden at concentrations that show minimal activity in standard broth (Fig. 3C vs 3F), indicating that the macrophage environment dramatically potentiates ARQ efficacy — consistent with our acidic RNS data (Fig. 3D-E). No obvious changes in per-well GAPDH levels were observed, demonstrating that ARQ at the indicated concentrations did not induce cytostasis or cytotoxicity in macrophages (Fig. 3F). Collectively, these results establish that ARQ exhibits anti-tuberculosis activity both in acidic RNS conditions and within the macrophage niche.

### ARQ synergizes with INH against Msm and Mtb

We further evaluated the potential of ARQ as a combination therapy with first-line Mtb drugs. Using Msm, we observed that ARQ significantly potentiated the activity of INH, reducing its MIC_90_ by approximately 7-fold (Fig. 4A). In contrast, no appreciable drug interaction was detected for rifampicin, ethambutol, or streptomycin (Fig. 4A). This synergistic effect with INH was further confirmed in a checkerboard assay, whereby non-inhibitory concentrations of ARQ (10-20 fold below its MIC_90_) substantially enhanced INH potency across a range of doses (Fig. 4B). Furthermore, the observed synergy is translatable to Mtb, where ARQ (4 μM) enhanced INH potency across all tested concentrations (Fig. 4C). Under standard culturing conditions, ARQ reduced the MIC_90_ of INH from 1.0 μM to 0.2 μM (Fig. 4D). Within infected macrophages, ARQ also markedly augmented INH-mediated bacterial clearance, reducing intracellular Mtb burden at different INH concentrations (Fig. S6). Collectively, these data demonstrate that ARQ synergizes specifically with INH against mycobacteria.

**Fig. 4.**
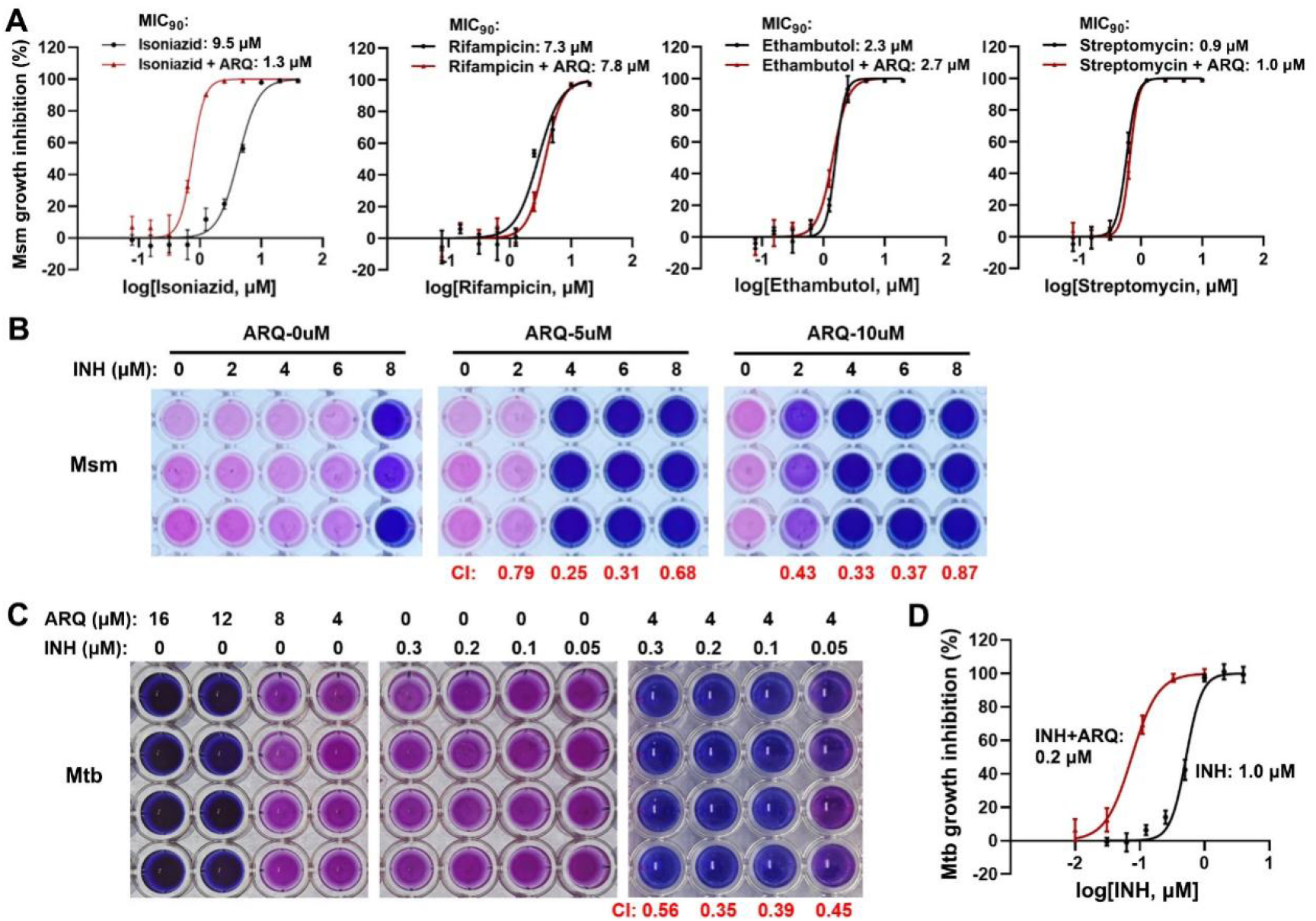
ARQ synergizes with INH against Msm and Mtb. **(A)** Dose-response curve of first-line drugs in the presence or absence of ARQ (10 μM) on Msm growth inhibition. **(B)** Resazurin microtiter assay (REMA) plates showing Msm growth in the presence of INH (0 - 8 μM) with ARQ at 0 μM, 5 μM, or 10 μM; combination indices (CI) are displayed below. **(C)** REMA plates for Mtb treated with different concentrations of ARQ and INH. **(D)** Dose-response curve of INH alone or INH + ARQ (3 μM) on Mtb growth inhibition, with MIC_90_ values indicated. Data are mean ± SD from at least triplicate experiments.

### KatG is a pupylation target that accumulates upon ARQ treatment

To elucidate the mechanism of ARQ’s synergy with INH, we performed a proteomic analysis of Msm treated with INH or INH+ARQ. The results revealed that ARQ treatment significantly upregulated 185 proteins and downregulated 225 proteins (Fig. 5A). Two proteases involved in protein quality control, Lon and Lon2, were upregulated 18-fold and 11-fold, respectively (Fig. 5A and Excel 1), consistent with the disruption of PafA-mediated protein homeostasis. Furthermore, many proteins involved in oxidative stress response, redox homeostasis, oxidoreductase activity, and disulfide reduction were upregulated, indicating that ARQ induced oxidative stress (Fig. 5B). Although ARQ has been reported to generate ROS in human cells via NQO1 targeting, this enzyme is absent in mycobacteria, and ARQ only induced ROS in the presence of INH (Fig. 5C). Therefore, the observed oxidative stress is likely attributable to enhanced INH activation, as the activated INH free radical is known to cause broad ROS stress beyond its primary target InhA.^20-22^

**Fig. 5.**
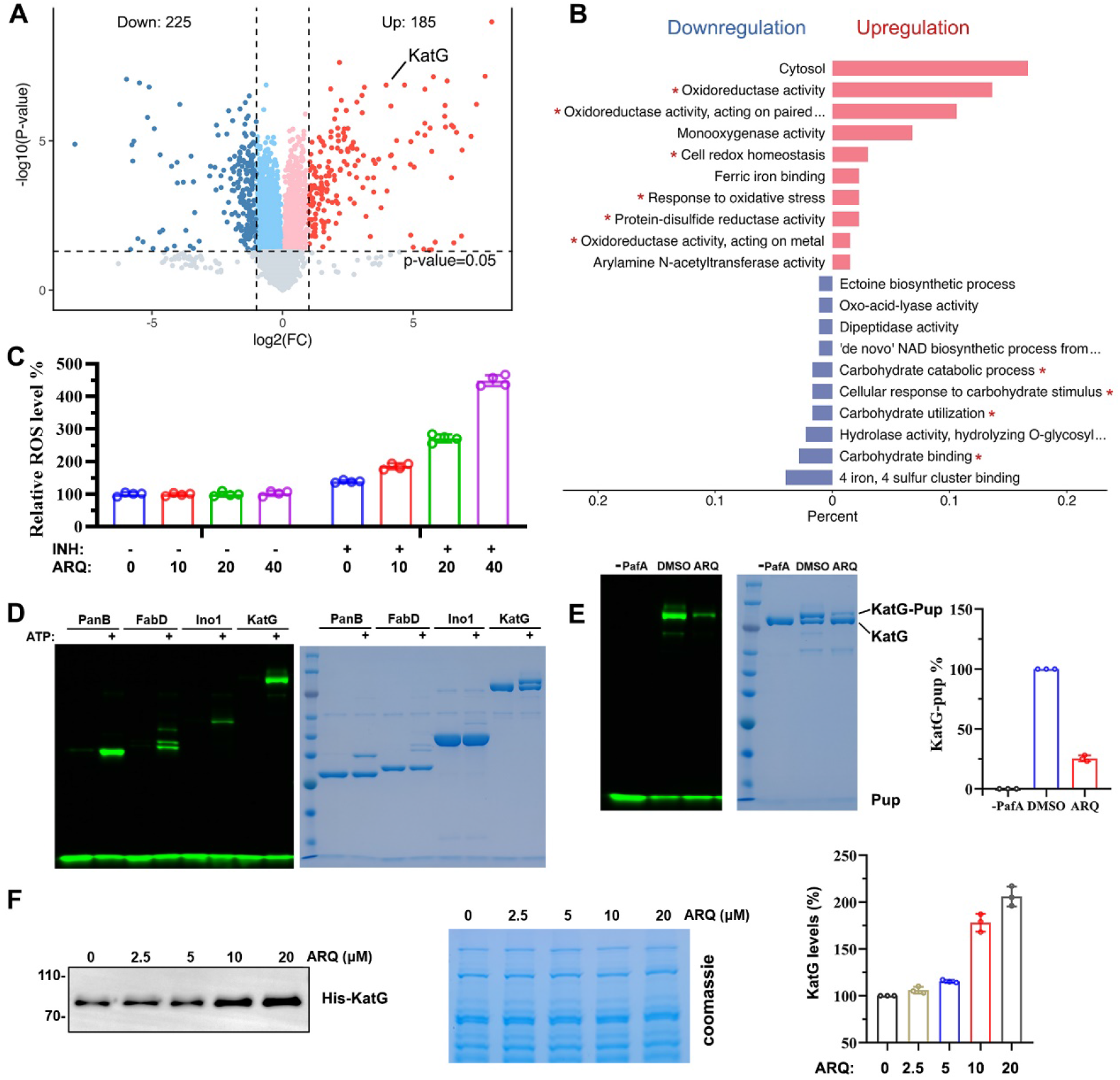
ARQ inhibits KatG pupylation to synergize with INH. **(A)** Volcano plot of proteomic changes in response to ARQ (comparing INH and INH+ARQ), with KatG highlighted. Protein level changes with |log_2_(FC)|>1 and p<0.05 are colored red (upregulated) and blue (downregulated). **(B)** Gene ontology analysis of differentially expressed proteins (comparing INH+ARQ vs. INH alone), highlighting molecular functions and biological processes affected by ARQ treatment. **(C)** Relative ROS levels in Msm treated with or without INH (2.5 μM, 24 h), showing that ARQ dose dependently increased ROS in the presence of INH. Mean ± SD, n=4. **(D)** Comparison of PafA-mediated pupylation (fluorescent and Coomassie-stained) of various substrates (PanB, FabD, Ino1, KatG) in the presence of ATP (200 μM). **(E)** ARQ (1 μM) inhibits PafA-mediated (0.2 μM) pupylation of KatG. **(F)** Level of ectopically expressed His-KatG in Msm lysates treated with increasing ARQ concentrations, detected using an anti-His antibody. The Coomassie-stained gel serves as loading control.

ARQ treatment was associated with changes in carbohydrate metabolism proteins (Fig. 5B and S7). Notably, we observed a 7-fold upregulation of KatG, the enzyme essential for activating the INH prodrug (Fig. 5A and Excel 1). We hypothesized that ARQ synergizes with INH by stabilizing KatG through the inhibition of its pupylation and degradation, thereby enhancing INH activation and amplifying oxidative stress. To test this, we first confirmed that KatG is modifiable by Pup, showing pupylation levels comparable to the known substrate PanB and exceeding those of FabD and Ino1 (Fig. 5D). Furthermore, ARQ markedly inhibited KatG pupylation in vitro (Fig. 5E), confirming that KatG is among the substrates whose pupylation is inhibited by ARQ. Consistent with our hypothesis, ARQ concentration-dependently increased the protein level of ectopically expressed His-KatG in *Msm* (Fig. 5F). This occurred in the absence of INH, and because ARQ alone does not elevate ROS (Fig. 5C), the increase in KatG levels cannot be attributed to ROS-mediated upregulation. Together, these results establish that ARQ inhibits KatG pupylation and degradation in mycobacteria, potentially explaining the observed synergy with INH.

### KatG accumulation drives ARQ-INH drug synergy and compensates for reduced KatG S315T activity in vitro

To establish the causal role of KatG regulation in drug synergy, we first constructed an anhydrotetracycline (ATC)-inducible KatG-depleted *Msm* strain using the plasmid pLJR962, which encodes both dCas9 and the gRNA. Results showed that ATC concentration-dependently abolished the activity of the ARQ-INH combination in *Msm* (Fig. 6A). At 5 μM ATC, the ARQ-INH drug synergy was completely lost (Fig. 6B). Conversely, KatG overexpression alone recapitulated a sevenfold synergistic effect with INH, mirroring the effect of ARQ treatment (Fig. 6C). As observed with depletion, ARQ conferred only minimal additional synergy in the strain already overexpressing KatG (Fig. 6C). These findings support a synergy model in which ARQ elevates KatG levels by inhibiting its PafA-mediated pupylation.

**Fig. 6.**
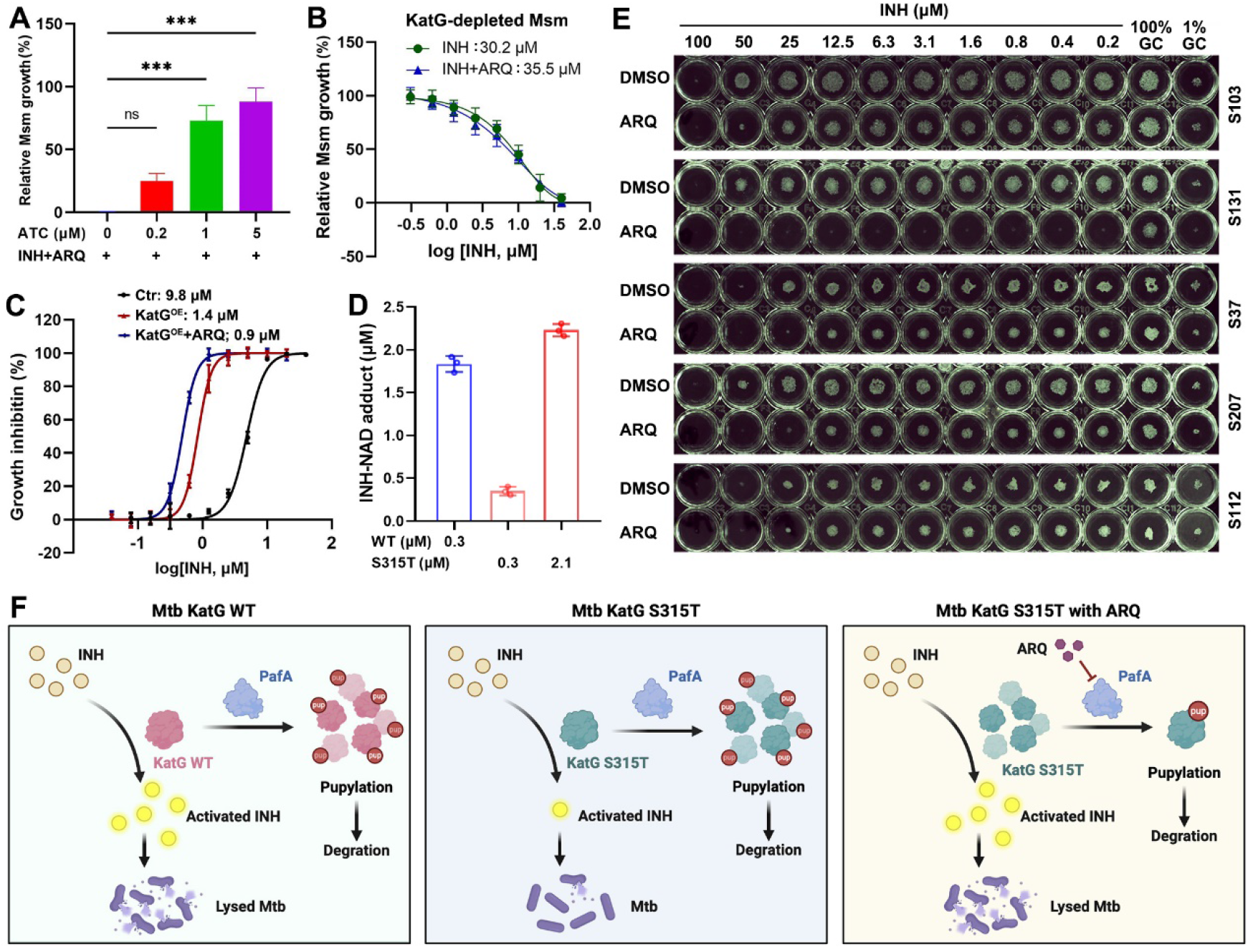
KatG accumulation drives ARQ-INH drug synergy and compensates for reduced KatG S315T activity in vitro. **(A)** ATC (anhydrotetracycline), which induces KatG depletion, dose-dependently reduced the anti-Msm activity of ARQ (10 μM) + INH (2.5 μM). **(B)** In KatG-depleted Msm (induced with 5 μM ATC), ARQ (10 μM) did not enhance the efficacy of INH. **(C)** Dose-response curve of INH alone or INH + ARQ against WT or KatG-overexpressing Msm, with MIC_90_ values indicated. Data are mean ± SD from at least three biological replicates. **(D)** INH-NAD adduct formation catalyzed by WT KatG and KatG S315T mutant. INH and NAD each 200 μM. Mean ± SD, n=3. **(E)** ARQ potentiates INH activity against clinical KatG S315T isolates to varying degrees. The 100% growth control (GC, no INH) tests whether ARQ alone has anti-Mtb activity. 1% GC = 1:100 dilution of the no-INH control. Colonies smaller than those in the 1% GC represent >99% growth inhibition. **(F)** Mechanism of ARQ-enhanced INH activity against mycobacteria. KatG is identified to be a pupylation target of PafA. ARQ inhibits PafA-mediated KatG pupylation, elevating KatG levels to boost INH activation. This enhances INH efficacy in wild-type Mtb and may overcome resistance in KatG-mutant strains.

Having established that ARQ acts by stabilizing KatG, we next asked whether this mechanism could overcome KatG-mutation-mediated INH resistance. The results showed that a sevenfold increase (mimicking the effect observed in our mass spectrometry data) in KatG S315T protein concentration completely compensated for the defect of INH activation in enzymatic assays (Fig. 6D). Therefore, ARQ-mediated KatG accumulation could, in principle, overcome INH resistance conferred by KatG mutations. To preliminarily test this, we evaluated five clinical INH-resistant *Mtb* isolates harboring the KatG S315T mutation. ARQ caused a >500-fold reduction in the INH MIC one of the strains (S131, Fig. 6E). The remaining four strains showed only 2- to 4-fold potentiation; however, ARQ notably reduced colony sizes in three of these strains across a broad range of INH concentrations (S103, S37, and S207). The basis for this variability is currently unknown but may include differences in baseline KatG expression levels, additional resistance mutations, or variations in PafA activity. A schematic model summarizing the mechanism of ARQ-enhanced INH activity is presented in Fig. 6F.

### ARQ is well-tolerated in mice over one week of treatment

Although BSL-3 constraints precluded efficacy testing in a mouse TB model, we assessed its potential toxicity, both alone and in combination with INH. A one-week treatment with ARQ alone (10 or 20 mg/kg) did not induce significant body weight loss (Fig. 7A). Co-administration of ARQ (20 mg/kg) with INH (10 or 20 mg/kg) was also well-tolerated (Fig. 7B). To assess potential organ toxicity, organ indices were calculated for the kidney, liver, lung, and spleen, which revealed no significant changes across all treatment groups, indicating a lack of gross pathological damage (Fig. 7C-F). Levels of serum creatinine (CREA-S, a marker of kidney function), as well as aspartate aminotransferase (AST) and alanine aminotransferase (ALT, markers of liver injury), remained within normal ranges across all treatment groups (Fig. 7G-I). Finally, histopathological examination of liver and kidney sections via H&E staining showed no obvious morphological abnormalities, even in the high-dose combination groups (Fig. 7J). Together, these data demonstrate that ARQ, alone or in combination with INH, is well-tolerated in mice over one week at doses up to 20 mg/kg, with no overt signs of acute toxicity.

**Fig. 7.**
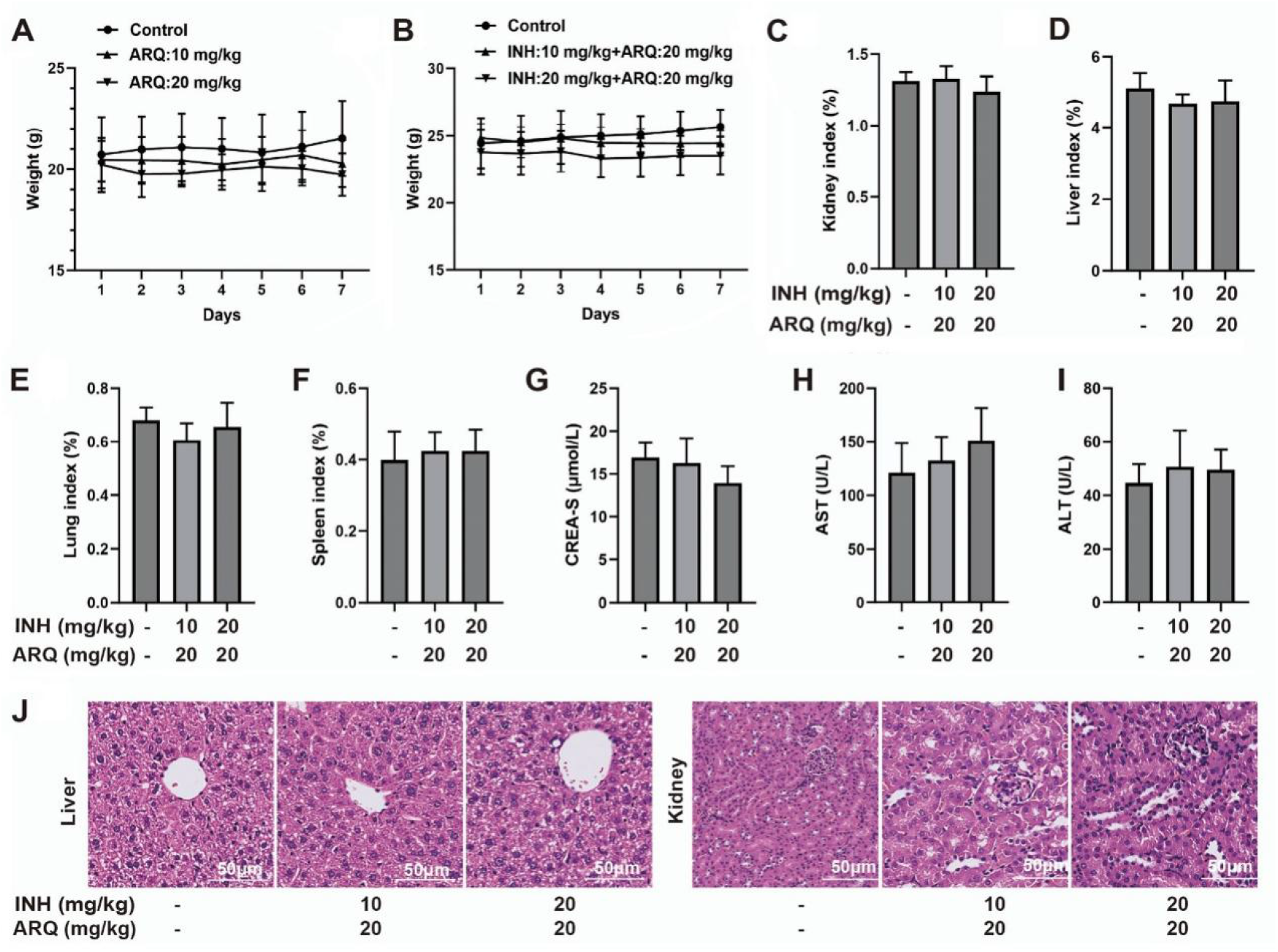
ARQ is well-tolerated in mice over one week of treatment. **(A)** Body weight changes over 7 days in mice treated with ARQ (10, 20 mg/kg) or vehicle control. **(B)** Body weight changes over 7 days in mice treated with INH (10, 20 mg/kg) combined with ARQ (20 mg/kg) or vehicle control. **(C-F)** Organ indices (kidney, liver, lung, spleen) calculated as (organ weight/body weight) × 100% for mice treated with INH (0, 10, 20 mg/kg) and ARQ (0, 20 mg/kg). **(G-I)** Serum creatinine (CREA-S), aspartate aminotransferase (AST), and alanine aminotransferase (ALT) levels in mice treated with INH (0, 10, 20 mg/kg) and ARQ (0, 20 mg/kg). **(J)** H&E-stained liver and kidney sections from mice treated with INH (0, 10, 20 mg/kg) and ARQ (0, 20 mg/kg). Data are mean ± SD from five samples.

## Discussion

Our study identifies ARQ as a potent, covalent, and substrate-competitive inhibitor of PafA. With an enzymatic IC_50_ in the sub-micromolar range, ARQ exhibits > 30-fold greater enzymatic inhibitory potency over previously reported inhibitors such as ST1926 and bithionol.^14^ ARQ is the first substrate-competitive PafA inhibitor that appears to bind at the periphery of the substrate-binding pocket, blocking Pup conjugation to different protein substrates with varying efficacy. However, the exact site of the substrate-binding pocket remains incompletely understood.^23, 24^ Two observations support an on-target mechanism: 1) the pronounced enhancement of ARQ’s activity under acidic RNS conditions and within macrophages mirrors the phenotype of PPS-deficient strains,^25-27^ and 2) the concordant inactivity of SF in both pupylation inhibition and bacterial suppression.

We further identified a specific synergy between ARQ and the frontline prodrug INH and elucidated its mechanistic basis. By identifying the catalase-peroxidase KatG—the enzyme responsible for INH activation—as a novel pupylation substrate, we show that ARQ accumulates KatG via PafA inhibition, thereby boosting INH activation and efficacy. This model is robustly supported by our proteomic data, direct in vitro pupylation assays, and KatG overexpression studies. These findings extend the functional repertoire of the PPS to include INH resistance, in addition to previously identified functions such as protein quality control, resistance to host-derived stress, and resistance to antifolates.^28, 29^

We further demonstrated that the reduction in catalytic activity caused by the prevalent KatG S315T mutation can be functionally compensated by increasing the concentration of the mutant enzyme. This “quantity over quality” approach relies on the residual enzyme activity of mutant KatG, analogous to strategies that optimize INH membrane trafficking.^30^ Consequently, it is not expected to work in strains that lack any KatG activity, including those with complete katG deletions.^31^ These findings suggest a potential strategy to overcome KatG-mediated INH resistance, the most common cause of INH resistance in clinical isolates.

This study has several limitations. First, the covalent binding site of ARQ on PafA and its binding mode have not been experimentally determined. We have initiated mass spectrometry and mutagenesis studies to map the site, but the precise residues modified remain to be identified. Second, while ARQ phenocopies PPS-deficient strains under acidic RNS conditions, definitive genetic validation using conditional *pafA*-knockout Mtb strains is needed to fully verify its on-target activity in live bacilli. Third, the molecular basis for the variable efficacy of ARQ-INH combination therapy against clinical KatG S315T isolates remains unknown. Fourth, although short-term mouse toxicity assessments revealed a favorable safety profile for ARQ, follow-up in vivo therapeutic efficacy investigations are required to advance preclinical development.

Despite these limitations, ARQ represents a significant advance in PafA-targeted drug discovery. It is the most potent PafA inhibitor reported to date, with robust activity in macrophages and under host-mimicking stress. The mechanism we have uncovered— stabilizing KatG via PafA inhibition to potentiate INH—offers a rational combination strategy to enhance frontline therapy and potentially overcome certain forms of INH resistance.

## Materials and Methods

### Plasmid construction, protein expression and purification

Mtb PafA-8T, Mtb PanB, Mtb KatG, Mtb FabD, and Mtb Ino1 were cloned into a pET-28a(+) expression vector containing an N-terminal 6×His-tag. Mtb Pup (39-64, Q64E) was synthesized and labeled with an N-terminal FITC fluorescent marker. The plasmids were transformed into E. coli BL21 (DE3) and cultured in LB medium. Protein expression was induced with 0.5 mM isopropyl β-D-1-thiogalactopyranoside (IPTG) for 3 h at 37 °C. Cells were harvested and lysed by sonication in a buffer containing 50 mM Tris-HCl (pH 8.0), 300 mM NaCl, 10% glycerol, and 1 mM PMSF. Proteins were purified using Ni-NTA beads, followed by Q-Sepharose anion-exchange chromatography (low-salt buffer: 20 mM Tris, pH 8.0, 10% glycerol, 1 mM DTT; high-salt buffer: 20 mM Tris, pH 8.0, 1 M NaCl, 10% glycerol, 1 mM DTT).

### Plate-based high-throughput screening assay

The assay was adapted from the previously established method with minor modifications. Briefly, PanB (20 μg/mL) was immobilized on a 96-well plate, followed by the addition of PafA (0.1 μM), FITC-Pup (1 μM), ATP (200 μM), and test compounds (2 μM). The samples were incubated at 37 °C for 30 min. After washing, fluorescence was measured using a Synergy H1 Hybrid Multi-Mode Reader (BioTek) at 480/520 nm.

### In-solution pupylation assay

Different substrates (4 μM), PafA (0.1 μM), FITC-Pup (4 μM), and ATP (200 μM) were incubated in reaction buffer (20 mM HEPES, pH 7.5, 300 mM NaCl, 10% glycerol, 2 mM DTT, 2 mM MgCl_2_). Reaction products were resolved by SDS-PAGE, and FITC signals were detected using a fluorescence imaging system (excitation: 488 nm). Total protein was assessed by Coomassie brilliant blue staining.

### In-Msm pupylation assay

Mtb Pup was cloned into a pMV261 expression vector (N-terminal HA-tag) and electroporated into M. smegmatis (Msm). Positive clones were cultured in LB medium with 5% glycerol and 0.1% Tween-20 (37 °C, shaking) until OD_600_ reached 0.6. HA-Pup expression was induced with 0.6% acetamide, and compounds were added simultaneously. After 3 days, cells were harvested, and HA-tagged proteins were detected by Western blot.

### Msm/Mtb MIC determination

The Msm pellet was collected at Log-phase and adjusted to OD_600_ = 0.002, inoculated into 96-well plates (100 μL/well). An equal volume (100 μL/well) of two-fold serially diluted compound solutions was added to each well. After 7-day incubation at 37°C, 0.01% resazurin indicator was added to each well. Following an additional 6-hour incubation, fluorescence intensity was measured at 590 nm (excitation 530 nm) using a microplate reader. For Mtb, the suspension was adjusted to OD_600_ = 0.008 and inoculated into 96-well plates (130 μL/well). To obtain measurable Mtb counts under acidic RNS stress, the initial inoculum was increased 25-fold (OD_600_ = 0.2). An equal volume (130 μL/well) of serially diluted compounds was added. After 14-day incubation at 37°C, 0.01% resazurin was added, and the plates were incubated for additional 24 hours. Minimum Inhibitory Concentration (MIC) values were determined using GraphPad Prism software.

### Acid RNS culturing of Msm/Mtb

Msm suspensions were prepared as Log-phase and diluted to OD_600_ = 0.005 using either pH 6.8 basal medium (7H9) or acidic medium (pH 5.5, adjusted with HCl) containing 3 mM NaNO_2_. Five milliliters of suspension was aliquoted into culture tubes. Test compounds (with DMSO controls) were added to the glass culture tubes. After 5-day shaking culture at 37°C, OD_600_ was measured. For Mtb, suspensions were prepared similarly and diluted to OD_600_ = 0.2 using either pH 6.8 basal medium or acidic medium with 3 mM NaNO_2_.

### Proteomics analysis

Msm cultures (OD_600_ = 1.0) were centrifuged at 200 × g for 10 min and diluted to OD_600_ = 0.5. The control group received 20 μM INH alone, while the experimental group received both 20 μM INH and 10 μM ARQ (triplicate biological replicates). After 24-hour shaking culture at 37°C, samples were collected by centrifugation (4°C, 3500 × g, 10 min), washed three times with pre-cooled PBS. Proteomic analysis was performed by OE Biotech (Shanghai, China) using DIA proteomics technology method.

### Survival in macrophages

THP-1 cells (50,000/well) in 12-well plates were treated with 100 nM PMA for 24 hours until adherence, then induced to M1 phenotype with 100 ng/ml LPS and 50 ng/ml IFN-γ for 36 hours. Mtb H37Ra strains were inoculated at an MOI of 5:1 (bacteria to cell ratio) and incubated for 4 h to allow phagocytosis; thereafter, each well was washed three times. Test compounds were added and cells were collected after 3 days. Following cell lysis, bacteria were diluted, plated, and cultured for 14 days before colony counting.

### Animal experiments

All animal studies were performed in accordance with the Ethical Guidelines for the Use and Care of Laboratory Animals and were approved by the Animal Ethics Committee of Sichuan Provincial People’s Hospital (approval number: 2025-660). C57BL/6 mice (18-20 g, 6 weeks, males) were obtained from SPF Biotechnology Co., Ltd. (Beijing, China), housed in a controlled environment (12 h light-dark cycle at room temperature 23±1°C) with free access to food and water. Mice were randomly assigned to three groups: control (Solvent), ARQ (20 mg/kg) group, INH (10 mg/kg) + ARQ (20 mg/kg) group, and INH (20 mg/kg) + ARQ (20 mg/kg) group. INH was dissolved in normal saline, while ARQ was dissolved in a solvent comprising 10% DMSO, 40% PEG-400, 5% Tween-80, and 45% normal saline. Intraperitoneal injections were administered daily for seven consecutive days. Mice were weighed, and the weights of the kidneys, liver, lungs, and spleen were recorded. Organ indices were calculated as the ratio of organ weight to body weight. Following anesthesia with avertin, blood was collected via the orbital sinus to measure biochemical parameters (including CREA-S, AST, and ALT). The liver and kidney tissues were fixed with 4% paraformaldehyde, dehydrated in a series of graded ethanol solutions, embedded in paraffin for slicing into 5 μm slides, and used for subsequent hematoxylin-eosin (H&E) staining.

## Acknowledgments

None.

## Author contributions

Conceptualization: CL, QS

Methodology: CL, BH, HX, QY, PX, TL, QS

Investigation: CL, BH, HX, QY, YL, CF

Visualization: CL, BH, QY, QS

Supervision: YL, PX, TL, QS

Writing-original draft: QS

Writing-review & editing: PX, TL, QS

## Data availability

All data are available in the main text or the supplementary materials.

